# Critical dynamics are a homeostatic set point of cortical networks *in vivo*

**DOI:** 10.1101/503243

**Authors:** Zhengyu Ma, Gina G. Turrigiano, Ralf Wessel, Keith B. Hengen

## Abstract

The brain is constantly challenged by severely destabilizing forces: proteins turn over rapidly, Hebbian modifications alter and introduce positive feedback into networks, and environments change over many timescales. Homeostatic plasticity mechanisms, which operate via negative feedback, are believed to compensate for these changes and constrain neuronal activity to a firing rate (FR) set point^1,2,3^. For decades, it has been widely assumed that activity in neural networks is robust as a direct result of the widespread expression of FR homeostasis^1,4^. Here we reveal that network dynamics are stabilized independent of excitatory FR homeostasis and that cortical networks actively self-organize around an ideal computational regime.

We continuously tracked network spiking activity in the visual cortex (V1) of freely behaving rats for nine days. We found that, under baseline conditions, networks of excitatory neurons are robustly organized around criticality, a regime known to maximize information capacity and dynamic range. Monocular deprivation (MD) revealed a dissociation of excitatory FRs and network dynamics. MD immediately and severely disrupted network organization, which returned precisely to criticality over 48h. In contrast, both the excitatory FR drop and the subsequent FR recovery trailed the timecourse of network changes by more than 30h. Model investigations suggest a role for inhibitory neurons in maintaining critical dynamics. Collectively, these results show that complex activity in cortical circuits is actively maintained near criticality and that this organization is not explained by previously identified mechanisms of pyramidal neuron FR homeostasis.

---

Neocortical neurons have a variety of cell-autonomous and local microcircuit mechanisms that can counterbalance perturbations in activity^5^. Consistent with this, in the visual cortex (V1), long-term visual deprivation initially suppresses neuronal firing^6,7^, which then exhibits a homeostatic rebound to baseline levels over multiple days^6^. FR rebound is exhibited at the level of individual regular spiking units (RSUs; presumptive pyramidal neurons)^3,6^, congruent with the expression of homeostatic plasticity mechanisms^2,8^. As such, there is a widely-held assumption that network stability is, from the standpoint of homeostasis, an inevitable effect of stable neurons^9,10^. However, it remains unknown a) whether network dynamics are subject to active stabilization and b) whether stabilization of individual neuronal activity gives rise to stable networks.

Neural networks can converge on or very near an emergent regime of population dynamics called “criticality”^11,12,13^. Criticality is a network state poised at the boundary between strongly and weakly coordinated population activity that maximizes information capacity (entropy) and transmission^14^. Measures of criticality are independent of FR and network size, making it a conceptually attractive candidate for a signature of stable networks capable of normal information processing. Despite this, Shew et al. (Ref. 14) point out that “… experiments on *in vivo* homeostasis have not yet been performed with attention to the possibility that criticality may be the end goal of homeostatic processes.”

In this context, we sought to answer three questions: first, to what extent is the cortex critical under normal conditions, second, if cortical activity reflects criticality, is this an epiphenomenon or a network set-point, and third, does network stability arise as a byproduct of underlying cellular set points? To answer these questions, we utilized MD as a direct homeostatic challenge to cortical activity *in vivo*. Brief MD (1–2d) results in widespread LTD across V1^15^ and a >50% suppression of cellular activity^3^. Prolonged MD reveals homeostatic increases in synaptic strengths and intrinsic excitability^16^, and a rebound of neuronal activity to baseline levels^3^. Recording network spiking continuously in this context allowed us to directly ask whether criticality is a set-point of network dynamics. We followed extracellular, single unit spiking for 200 h in deprived and control hemispheres of monocular V1 (V1_m_) in freely behaving rat young rats (for details, see Ref. 3) and measured network state with respect to criticality. RSUs^6,17^ were classified as continuously recorded if spikes within the cluster were present for at least 80% of the 9-day recording period^3^ (46 RSUs from 7 animals). Units present for less than 80% of the recordings were classified as non-continuous and considered as individual units within each applicable 4h bin (MD: 10,280, Control: 8,584 RSUs from 7 animals). Measurements of network dynamics include all single units detectable in each 4h bin, while measurements of FR homeostasis are based on continuously observable units.

Critical systems are defined by scale free dynamics, such that events spanning all spatial and temporal scales are observed according to power laws. In this context, events are contiguous cascades of spiking activity^13,18^ colloquially termed “neuronal avalanches”. Avalanches were analyzed in terms of size (**S**, the number of spikes), and duration (**D**, time) (**Fig 1A**), and power law exponents were fit to the two distributions. In critical systems, the exponents of the two distributions can be used to predict the mean avalanche size (<S>) observed at a given duration (i.e. the distributions scale together)^19^. When <S> is plotted against avalanche duration, the difference between the empirically derived best-fit exponent and the predicted exponent serves as a compact measure of the deviation from criticality (**D**eviation from **C**riticality **C**oefficient, “DCC”, **Fig 1B**). This term, DCC, effectively identifies synthetic data tuned to subcritical, critical, and supercritical network states (**Fig 1C**). In control conditions, *in vivo* cortical circuits were organized close to criticality (DCC<0.2; **Fig 1D**).

**Figure 1.**
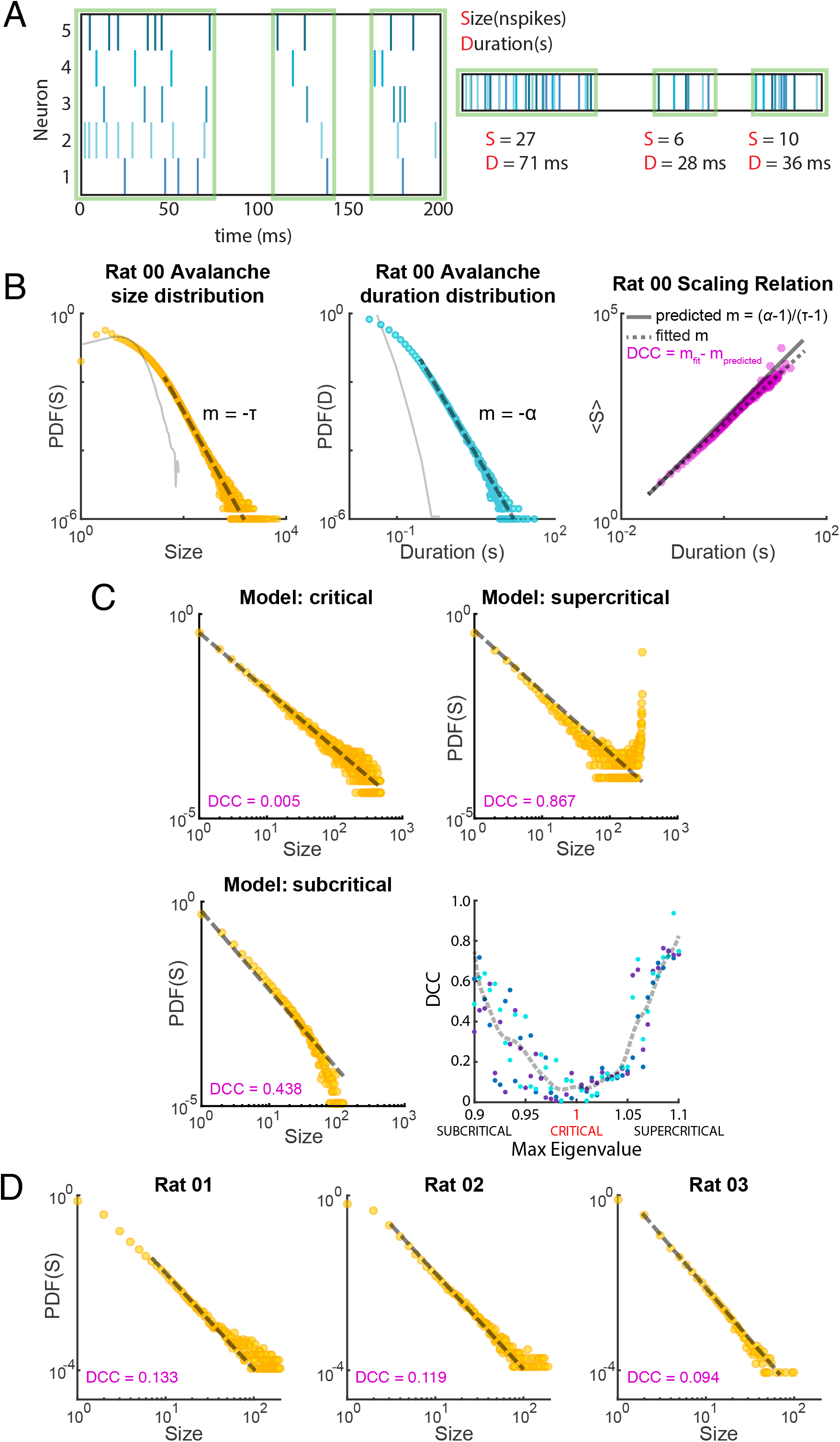
DCC, the Deviation from Criticality Coefficient, is an effective, scalar measure of how near a neural network is to criticality. All recordings were of regular spiking single units in V1 of freely behaving rats. (**A**) Discrete relaxation events, i.e. avalanches, are identified in ensemble recordings by the presence of silent periods. Spikes from all neurons in a region of interest can contribute to an avalanche, which is measured as a function of the number of contributing spikes (S) or the event duration (D). (**B**) (left, center) The probability (PDF, probability distribution function) of observing an avalanche of a given size (gold) or duration (blue) can be fit by power laws, each generating an exponent (–τ, –α). Solid gray traces display avalanche distributions derived from shuffled data. (right) In order for a system to be considered critical, the average avalanche size (<S>) at a given duration (purple) must scale according to a predicted fit (predicted exponent = (α−1)/(τ−1)). The difference between the predicted exponent (solid gray line) and the observed exponent (dashed gray line) can be used as a quantitative measure of nearness to criticality (Near Criticality Coefficient, DCC). (**C**) The avalanche size PDF and associated DCC extracted from model networks operating in critical (top left), supercritical (top right), and subcritical regimes (bottom left). (bottom right) In three model networks (three colors), the activity of every neuron is known at each timestep. Networks with a maximum eigenvalue of 1.0 are in a critical regime by definition. This known quantity is compared to DCC, which is derived from subsampling the network, a parallel to biological methods. (**D**) Visual cortical circuits operate near criticality as demonstrated by avalanche size PDFs and associated DCCs extracted from 4h of single unit data in each of three freely behaving rat pups. Data in B and D are derived from ensembles of well-isolated RS units. Model and empirical data have been fit with power law functions.

The combination of new recordings with earlier datasets further supported previous findings^3^. After 28h of MD, RSU spontaneous single-neuron FR was suppressed by ∼60%. By MD5 the same neurons exhibited a homeostatic rebound to baseline FRs (**Fig 2A**). These results mirror the time course of cellular responses to MD via lid-suture^6,15,16,20^. In stark contrast, cortical circuits deviated from the critical network state immediately upon light exposure following MD (*DCC ≈* 0.6, **Fig 2B**), more than 24h before FRs were perturbed. Cortical circuits then returned to near-criticality at the beginning of MD3 while FRs were at the nadir of their suppression. The rapid deviation from criticality on MD1 preceded a reduction in single-neuron FRs by nearly 30h (**Fig 2A,B**). Similarly, single-neuron FR homeostasis lagged the resurrection of network criticality by >32h (**Fig 2A,B**; individual animal data in **Fig S1,S2**). We note that control hemisphere FRs and network dynamics were unaffected by MD (**Fig 2C,D**). In addition, under baseline conditions (i.e. prior to lid suture), the network state in V1 operated near the critical regime (*DCC* ≈ 0.2) in both light and dark (**Fig 2B**). These data demonstrate that the near-critical network state serves as a central attractor of circuit dynamics in V1 of freely behaving animals. The timing of the deviation from the critical network state and the recovery towards thereof reveals that stable cortical network dynamics can emerge independently of excitatory single-neuron FRs and the biophysical elements of excitatory single-neuron FR homeostasis.

**Figure 2.**
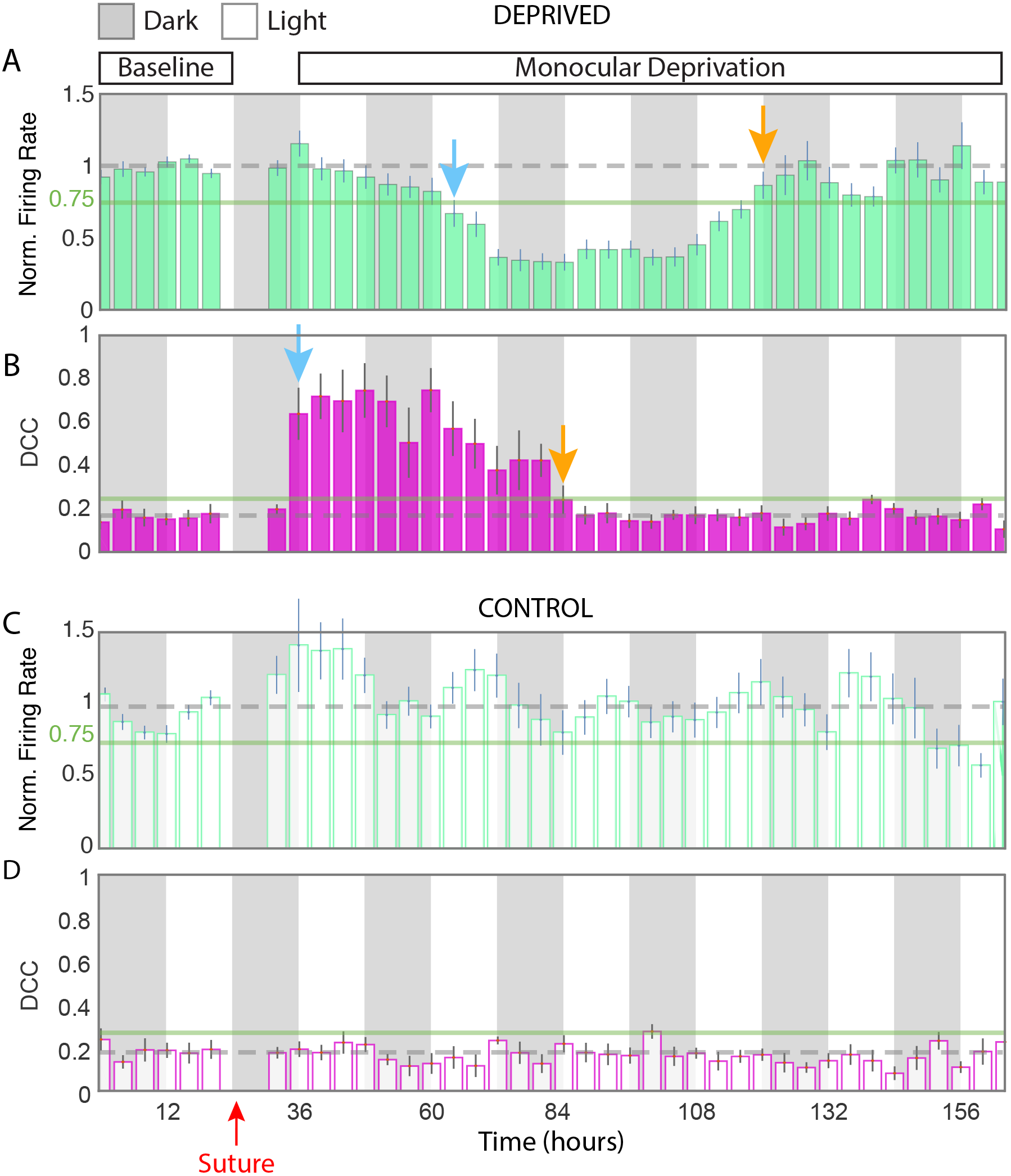
Network dynamics are homeostatically tuned to near-criticality, independently of excitatory FR homeostasis. (**A**) The FRs of continuously observable excitatory neurons followed across 200h recordings show a biphasic response to monocular deprivation (MD, 47 units, 7 animals). FRs were normalized to 24h of baseline recordings (gray dashed line) prior to the induction of MD. FRs were within 25% of baseline levels (green solid line) for more than 24h after the start (lights-on) of the first day of MD (MD1). Firing rate suppression past 25% is denoted by the blue arrow. FRs were maximally suppressed at 84 h and slowly rebounded to baseline levels, re-crossing the 25% line at 120 h (MD4), denoted by gold arrow. (**B**) Network dynamics were examined in the same recordings for nearness to criticality. Immediately upon light exposure on MD1, the mean DCC more than tripled, crossing the 25% line in the first bin (blue arrow). The mean DCC was restored to baseline levels at 84 h (MD3, gold arrow). (**C**) In the control (contralateral) hemisphere of animals contributing data to (A) and (B), MD had no significant impact on mean normalized firing rate of continuously observable units. (**D**) MD had no significant impact on mean DCC in control hemisphere recordings. FR/DCC data from the night before MD1 are not shown as they are subject to artifacts from the process of brief anesthesia and lid suture, and are thus not considered in calculation of baseline parameters.

We next sought to understand what aspects of cortical networks might support FR homeostasis and the emergence of an attractor near criticality. We created an abstract model whose key features resembled those found in V1 (**Fig S3**). Parameters were chosen based on empirically determined values, e.g. inhibitory neurons were the minority of neurons^21^, and were more broadly connected throughout the network than were excitatory neurons^22^. Both excitatory and inhibitory neurons expressed homeostatic (synaptic scaling; SS) and spike timing dependent (STDP, i.e. Hebbian) plasticity mechanisms. Briefly, SS was a global, multiplicative adjustment of all synapses onto a neuron that served to compensate for deviations in output. STDP was an activity dependent increase or decrease in the strength of a specific synapse. We subjected models to a sustained reduction of excitatory input (mimicking MD) and searched for subsets of model parameter combinations that replicated our empirical results, specifically 1) a deviation of the network state from criticality upon reduced input, 2) a reduction in FR, first of inhibitory neurons followed by excitatory neurons^6^, and 3) a homeostatic recovery of FR and criticality. We explored >400 combinations of three parameters: i) the fraction of inhibitory neurons in the network, ii) the fraction of excitatory neurons contacted by each inhibitory neuron, and iii) the fraction of inhibitory neurons receiving external input. Less than 0.5% of the models satisfied these constraints (**Fig 3A-D, S3**), suggesting that real cortical networks have selected for precise excitatory/inhibitory connectivity rules^23,24,25^. Neither manipulation of plasticity strength (10 to 150-fold change in gain, SS and STDP) nor excitatory to excitatory connectivity (1 to 30 percent) were capable of rescuing a failure regime adjacent to a successful topology (**Fig S3**). These results implicate precise inhibitory architectures in establishing stable cortical dynamics.

**Figure 3.**
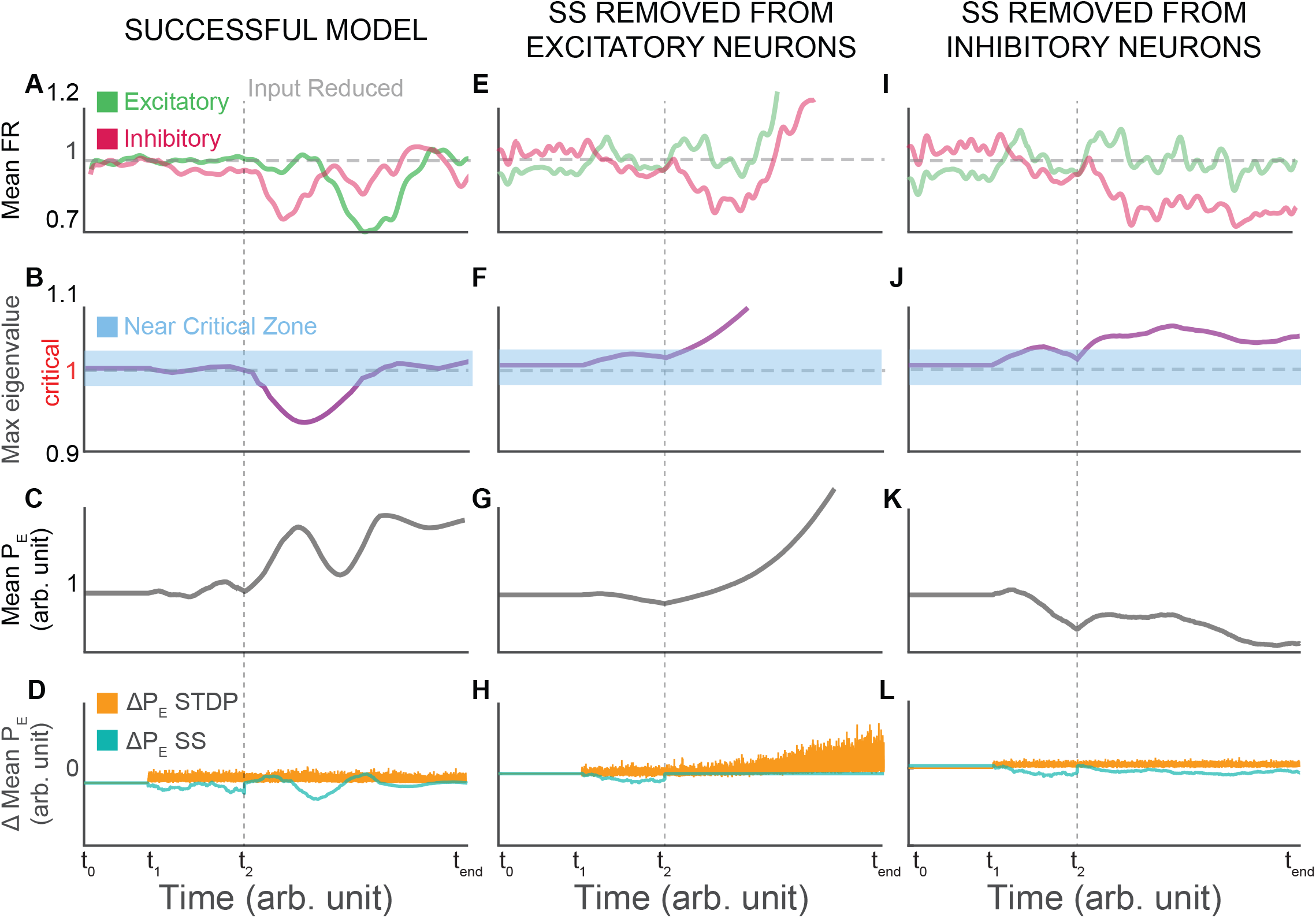
Model cortical networks composed of inhibitory and excitatory neurons were subjected to stable input for 20,000 simulation time steps (t_0_ to t_1_). Spike timing dependent plasticity (STDP) and synaptic scaling (SS; a global multiplicative compensatory change in synaptic strength) were turned on at 50,000 steps (t_1_). External input to the network (vertical dashed line, t_2_) was reduced as a homeostatic challenge mimicking monocular deprivation. (**A through D**) Successful models recapitulated empirical results, such that input reduction: suppressed FRs of inhibitory neurons (red) prior to excitatory neurons (green), and both rebounded to baseline levels by t_end_(**A**), and eliminated the critical network state (max eigenvalue equals 1) which rebounded by t_end_ (**B**). The mean excitatory synaptic strength (P) across the timecourse of successful models revealed a net increase by t_end_ (**C**). The progression of mean P as a function of changes in P due to STDP (orange) and SS (green) (**D**). (**E through H**) Successful models were rerun and SS was removed from excitatory neurons at the onset of input reduction. Inhibitory and excitatory FRs exhibited runaway gain (**E**) and network dynamics became unboundedly supercritical (**F**). Mean P exhibited a similarly unbounded progressive increase (**G**) as a result of uncompensated STDP (**H**). (**I through L**) Successful models were rerun and SS was removed from inhibitory neurons at the onset of input reduction. Neither excitatory nor inhibitory FRs exhibited runaway gain (**I**). Near critical network dynamics were eliminated (**J**). Mean P exhibited a net reduction (**K**) and alongside stable SS and STDP (**L**).

To probe the contributions of Hebbian and homeostatic plasticity to stable networks, we reran successful models and systematically turned off either SS or STDP in either excitatory or inhibitory neurons upon the initiation of input reduction. As expected, the deletion of SS from excitatory neurons resulted in runaway gain and severely destabilized FRs and network state^9,26^(**Fig 3E-H**). Elimination of STDP from either excitatory or inhibitory neurons largely eliminated the impact of input reduction all together (**Fig S4**). Remarkably, elimination of SS from inhibitory neurons pushed network dynamics out of the near critical state without destabilizing FRs (**Fig 3I-L**). Together, these model investigations make the strong prediction that homeostatic mechanisms in excitatory neurons are necessary for single-neuron control of FRs while stable circuit dynamics depend on the widespread expression of homeostatic mechanisms in inhibitory neurons.

The challenge of maintaining stability in neural circuits has been recognized for more than 40 years^1,27,28^. Efforts to understand this process have revealed homeostatic mechanisms at the level of ion channels, synapses, membrane excitability, and FRs^1,6,16,29^. However, the stability of neural circuits and networks, especially in the intact brain, has remained addressed only by computational and theoretical works^15,26^. Here we provide a direct demonstration of circuit-level stabilization in the intact brain that does not depend on previously identified mechanisms. These findings implicate criticality as a central attractor for cortical organization^30^ and reveal that cortical circuits can self-organize around a computationally idealized regime despite ongoing suppression of spiking activity. That the critical network state in visual cortex spans extended periods of time as well as transitions in light and dark suggests that it is not a result of a subset of initial conditions but a generalized rule. Our model investigations suggest that inhibitory architecture and regulation is essential for the establishment of a topology capable of supporting criticality. Studying the contribution of cell types, wiring patterns, and plasticity mechanisms to complex network dynamics will be essential to understand the robustness of neural computation and long-term information processing in the brain.

