## Supplemental Figures and Methods for "Critical dynamics are a homeostatic set point of cortical networks *in vivo*"

### FIGURE S1

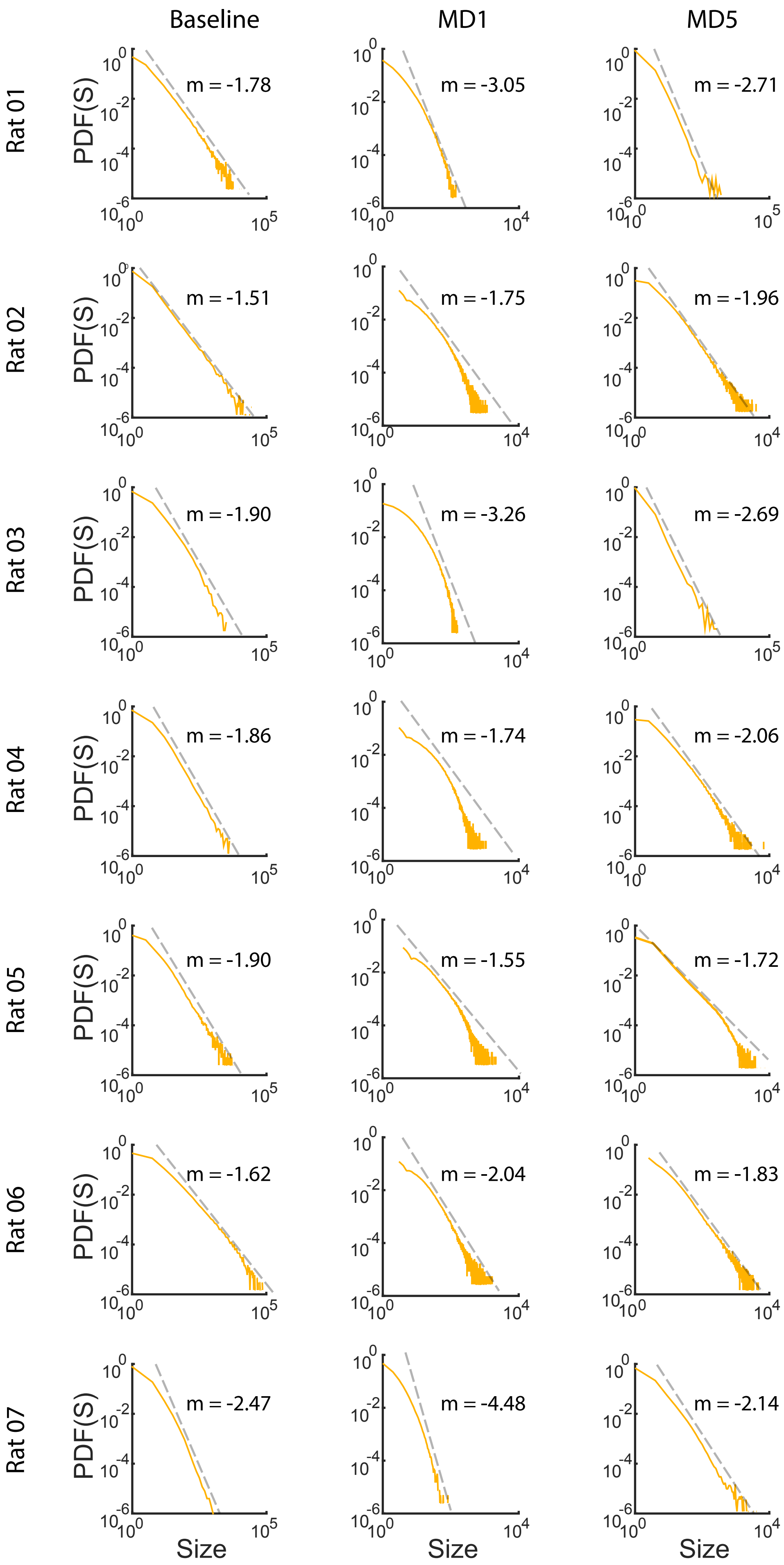

Baseline
MD1
MD5
Rat 01
Rat 02
Rat 03
Rat 04
Rat 05
Rat 06
Rat 07
PDF(S)
Size
m = -1.78
m = -3.05
m = -2.71
m = -1.51
m = -1.75
m = -1.96
m = -1.90
m = -3.26
m = -2.69
m = -1.86
m = -1.74
m = -2.06
m = -1.90
m = -1.55
m = -1.72
m = -1.62
m = -2.04
m = -1.83
m = -2.47
m = -4.48
m = -2.14

FIGURE S2

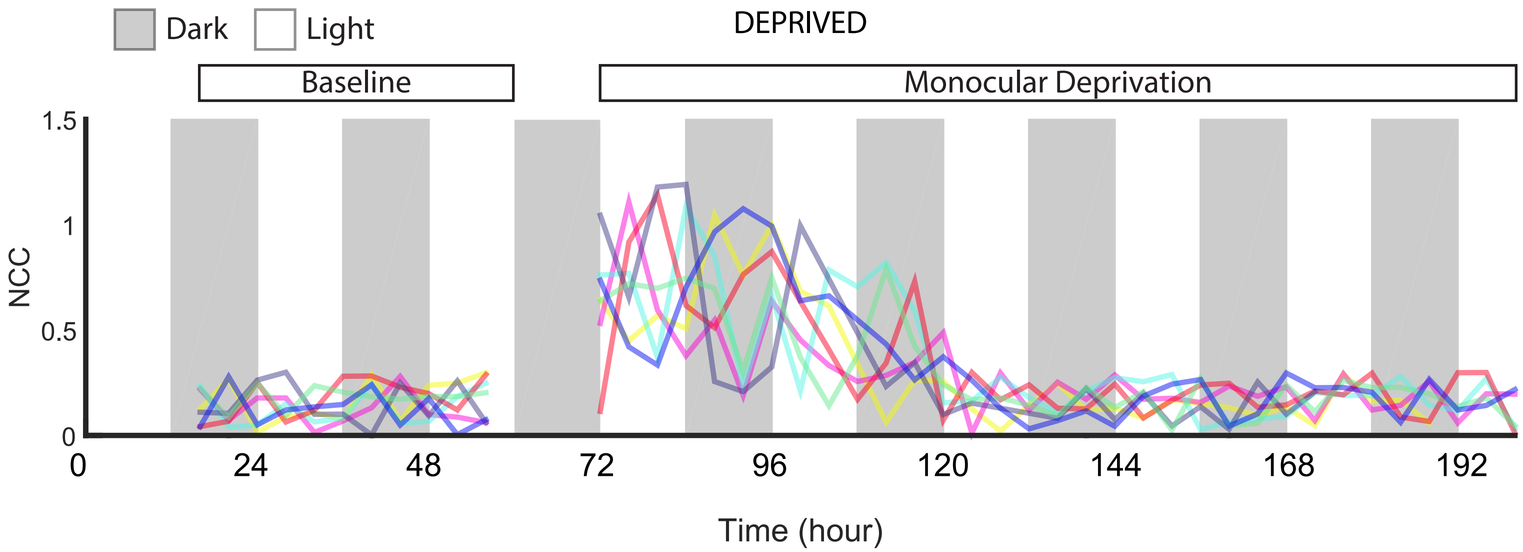
Dark
Light
DEPRIVED
Baseline
Monocular Deprivation
1.5
1
0.5
0
NCC
0
24
48
72
96
120
144
168
192
Time (hour)

FIGURE S3

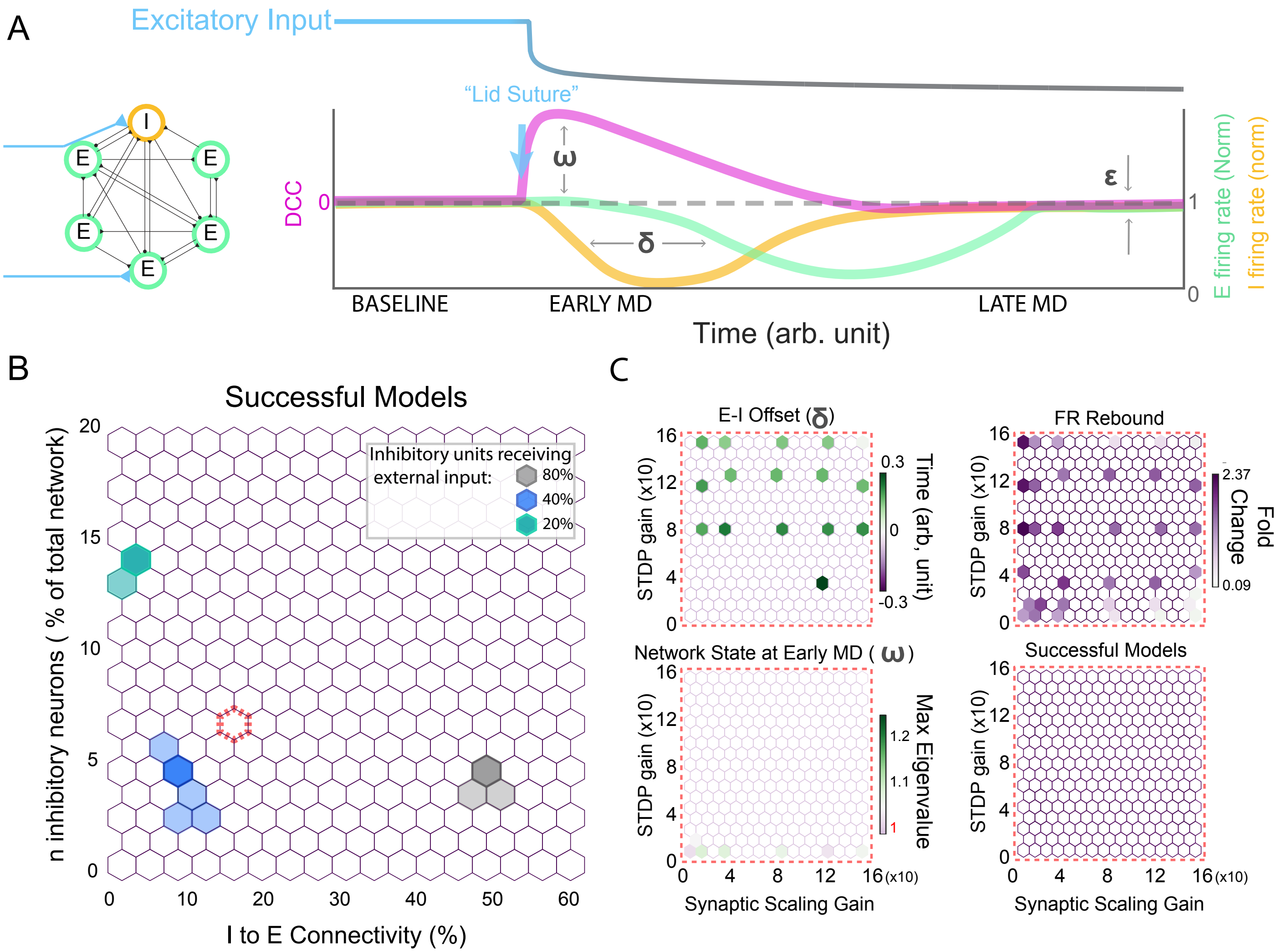

A
Excitatory Input
"Lid Suture"
I
E
E
E
E
E
DCC
0
ω
ε
δ
1
0
E firing rate (Norm)
I firing rate (norm)
BASELINE
EARLY MD
LATE MD
Time (arb. unit)
B
Successful Models
Inhibitory units receiving external input:
80%
40%
20%
n inhibitory neurons ( % of total network)
I to E Connectivity (%)
C
E-I Offset (δ)
Time (arb, unit)
FR Rebound
Fold Change
2.37
0.09
Network State at Early MD ( ω )
Max Eigenvalue
Successful Models
STDP gain (x10)
Synaptic Scaling Gain

FIGURE S4

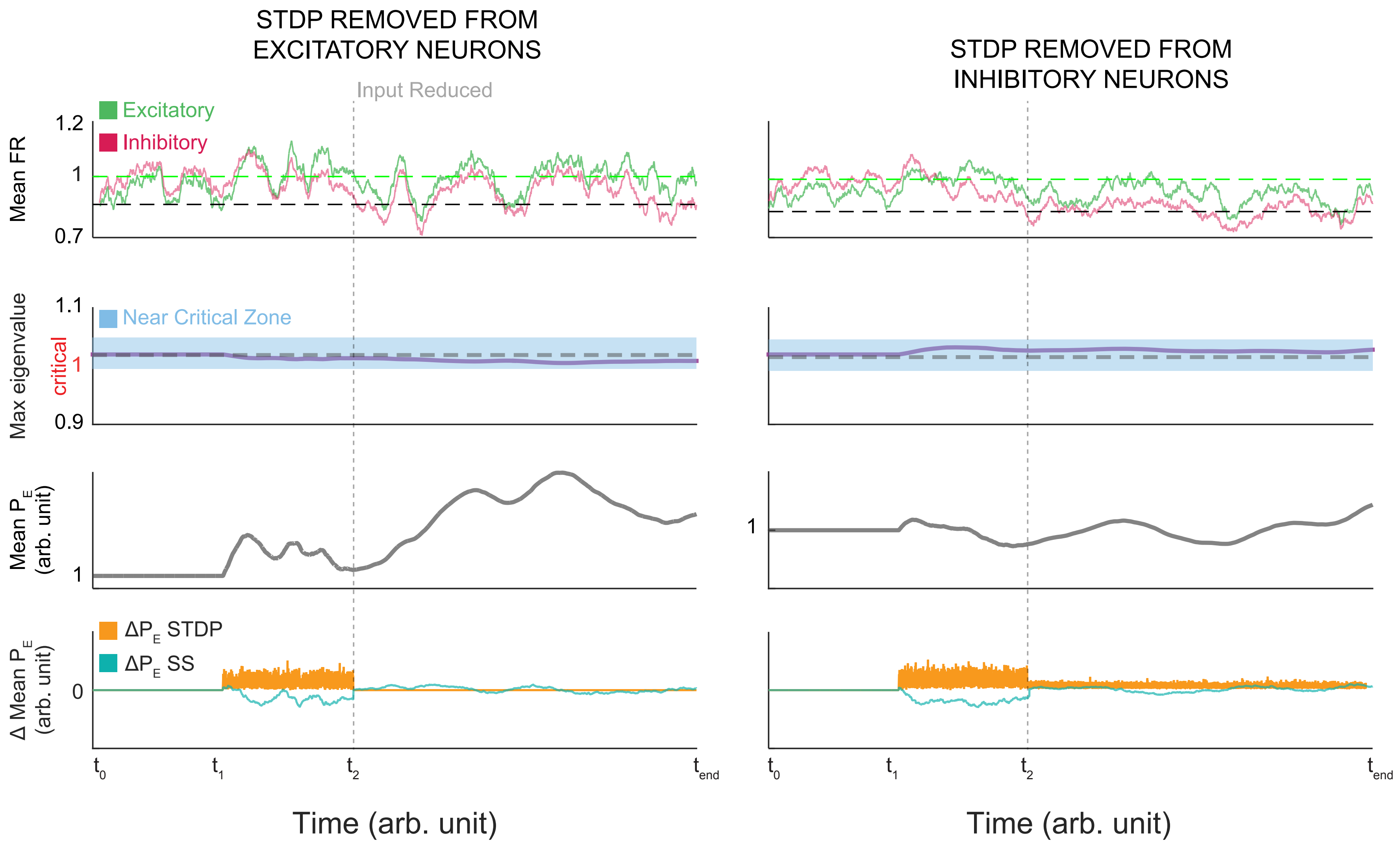

STDP REMOVED FROM EXCITATORY NEURONS
STDP REMOVED FROM INHIBITORY NEURONS
Input Reduced
Excitatory
Inhibitory
Mean FR
1.2
1
0.7
Near Critical Zone
Max eigenvalue
critical
1.1
1
0.9
Mean $P_E$ (arb. unit)
1
$\Delta P_E$ STDP
$\Delta P_E$ SS
$\Delta$ Mean $P_E$ (arb. unit)
0
$t_0$
$t_1$
$t_2$
$t_{end}$
Time (arb. unit)

**Figure S1** Avalanche size PDF plots from all animals at baseline, early MD, and late MD reveal a consistent shift into subcritical dynamics in early MD followed by a return to power law distributions. Avalanches are composed of spatiotemporally clustered bursts of action potentials generated by well-isolated RS units in monocular primary visual cortex. The probability of observing an avalanche of a given size (gold) was with a single exponent (power law; m). Avalanche distributions are shown for all seven freely behaving animals used in this study, at three key time points. Baseline refers to pre-monocular deprivation, MD1 refers to the first day of monocular deprivation when firing rates (FR) of the constituent single units was still at baseline levels, and MD5 refers to the fifth day of monocular deprivation when both FRs and DCCs (near criticality coefficient) displayed homeostatic recovery.

**Figure S2** Individual animal DCC traces from the deprived hemisphere. Network dynamics were examined in the each of seven recorded animals for nearness to criticality (near criticality coefficient, DCC). In seven of seven animals, light exposure on MD1 caused the mean DCC to increase significantly. The mean DCC in all animals recovered to baseline levels between 120 and 124h. DCC data from the night before MD1 are not shown as they are subject to artifacts from the process of brief anesthesia and lid suture, and are thus not considered in calculation of baseline parameters.

**Figure S3** Inhibitory parameters and not excitatory parameters or synaptic plasticity rules are sufficient to recapitulate empirical results in a series of models. (**A**) (left) Illustration of model recurrent network with excitatory input (blue) to inhibitory (orange) and excitatory (green) neurons. (right) Illustration of model time course and “successful” results. Firing rates of excitatory (E) and inhibitory (I) neurons and network state with respect to criticality (DCC) were monitored continuously. In successful models, “lid suture” (reduction in excitatory input, blue/gray line) rapidly increased DCC ($\omega$), suppressed inhibitory neuron FRs prior to suppressing excitatory neuron FRs ($\delta$), and each of these measures returned to baseline parameters ($\varepsilon$) despite maintained reduction in input. (**B**) The inhibitory fraction of the network and the number of excitatory neurons contacted by each inhibitory neuron were systematically varied across three levels of input to inhibitory neurons. Only three discrete combinations of parameters were sufficient to reproduce empirical results (green, blue, and gray). (**C**) A reasonable (nearby) but unsuccessful arrangement of inhibitory parameters was selected (dashed red line in (B)). With these parameters fixed, no explored region in the three-dimensional space defined by homeostatic plasticity gain (synaptic scaling, SS), spike timing dependent plasticity gain (STDP), and excitatory to excitatory neuron connectivity (%) was capable of rescuing the model. STDP gain factor was applied to both inhibitory and excitatory terms.

**Figure S4** Model cortical networks composed of inhibitory and excitatory neurons were subjected to stable input for 20,000 simulation time steps ($t_0$ to $t_1$). Spike timing dependent plasticity (STDP) and synaptic scaling (SS; a global multiplicative compensatory change in synaptic strength) were turned on at 50,000 steps ($t_1$). External input to the network (vertical dashed line, $t_2$) was reduced as a homeostatic challenge mimicking monocular deprivation. Successful models recapitulated empirical results (see Fig 3). Successful models were rerun and STDP was removed from excitatory neurons (left column) or inhibitory neurons (right column) at the onset of input reduction. In neither case were FRs severely perturbed (top) and in neither case did network dynamics ever exit the near critical zone (middle). Mean excitatory synaptic strength (P; third row) and underlying changes in P as a function of STDP and SS are shown (bottom row).

**SUPPLEMENTAL**

**Methods:**

All surgical techniques and procedures were conducted in accordance with the Brandeis University IACUC and NIH guidelines.

**Previous Datasets**

We combined two new recordings (see below) with previously published data from five animals (Hengen et al., 2016).

***In Vivo* Data Acquisition**

Experimental design, extracellular recordings, and data collection were as described previously[1,2]. Briefly, on postnatal day 21 (P21), Long-Evans rat pups of both sexes were anesthetized and a small craniotomy was drilled over monocular primary visual cortex of each hemisphere. Dura was resected and 16ch microelectrode arrays (Tucker-Davis Technologies) were stereotaxically placed with wires spanning all 6 cortical layers. Single wires were implanted bilaterally into the nuchal muscles for electromyographic signal (EMG) acquisition. Animals were allowed to recover from surgery for two days. On the second day of recovery, animals were placed in the environmentally enriched recording chamber (which replicated a home-cage environment) with a litter mate. Continuous data acquisition began after lights-on (ZT0) on P24. Baseline activity was recorded for 24h (ZT12 of P25 to ZT12 of P26). After lights-out (ZT12) at the end of baseline, animals were briefly disconnected (5-15 minutes) from recording tethers and one eye was sutured. Animals were promptly returned to the recording chamber in complete darkness. Due to the neuronal impact of brief anesthetization, data from the impacted bins of the baseline (P26 dark period) were ignored. The commencement of monocular deprivation (MD) was considered lights-on (ZT0) of P27, at which time animals experienced stimulus-driven ocular disparity for the first time. Recordings were maintained continuously for the next 132h (MD1-MD6). Animal behavior was recorded via synchronized video. All neural data were sampled at 30 kHz and broadband signals were written to disk for offline processing.

**Data Processing**

Neural data were bandpass filtered (300 to 10,000 Hz) and thresholded for spike detection (-3.5 SD). Spikes were interpolated, spline fit, and the first four principal components (PCs) of the waveform matrix were calculated. Spike PCs were clustered with Klustakwik[3]. Clusters generated by single units were separated from those arising from multiunits with a 15-node random forest trained on >2,000 single units from previous recordings. The most important feature in single-unit detection was the degree of refractory period contamination. Tested against manually scored datasets, this automated processing generated over 93% agreement. All machine-scored datasets were subsequently manually checked.

Cells were considered continuous if 1) clusters did not drift, 2) single-unit properties were maintained from the beginning of the baseline period until hour 134 (80% of the recording), 3) biophysical properties were consistent with single units (e.g. refractory period, stable waveform), and 4) Signal to noise ratio was high throughout the period considered (>=80% of the recording). Single units that were not detectable for at least 80% of the recording were stored separately as transient units. Regular spiking units (RSUs) and fast spiking units were identified based on waveform shape[4,5]. Only RSUs were considered in this work. The baseline recording (24h) before lid-suture was used for firing rate normalization of continuous units.

**Avalanche Analysis**

We discretized time into 40 ms bins, binarized each neuron's spike train into its "activity", i.e., 0 or 1, and obtained the "network activity" as the sum of all recorded activity within the time bin. Based on the network activity, we defined a "neuronal avalanche" by introducing a threshold at 35 percentile of total network activity[6,7]. An avalanche starts when the network activity crosses the threshold from below and ends when the network activity crosses the threshold from above. We quantified each neuronal avalanche by its size S, i.e., the integrated network activity between threshold crossings, and its duration D, i.e., the time between threshold crossings. Using maximum likelihood estimation methods, we fitted a truncated power law $f(S) = \frac{S^{-\tau}}{\sum_{Smin}^{Smax} S^{-\tau}}$ to the avalanche size distribution of $N_{av}$ avalanches using the following iterative procedure. (i) The maximum avalanche size $S_{max}$ was taken as the largest observed avalanche size. (ii) The exponent $\tau$ was estimated for three values of the minimum avalanche size $S_{min}$ ranging from 1 to 10 and the corresponding Kolmogorov-Smirnov (KS) values were obtained. (iii) The minimum avalanche size $S_{min}$ and the corresponding exponent $\tau$ yielding the smallest KS value were chosen. (iv) When $KS < 1/\sqrt{N_{av}}$, the exponent estimation was completed. Otherwise, the procedure (ii) to (iv) was repeated with the maximum avalanche size $S_{max}$ reduced by 1 until the condition $KS < 1/\sqrt{N_{av}}$, was satisfied. We applied the same fitting procedure to the avalanche duration distributions with corresponding exponent $\alpha$. To evaluate whether a power law was a plausible fit of an avalanche distribution, we performed hypothesis testing. We simulated 1000 artificial power law distributions (surrogate distributions) with the same exponent, number of avalanches, minimum avalanche size, and maximum avalanche size, as estimated from the experimental avalanche distribution. Specifically, using the inverse method, the surrogate distributions were generated according to $S = S_{min}(1 - r)^{-1/(\tau-1)}$ where $r$ was a random number drawn from a uniform distribution between 0 and 1. Thereafter, the distribution was upper-truncated by setting a cut-off at the maximum value $S_{max}$ observed in the empirical data. The deviation between the simulated surrogate distributions and a perfect power law was quantified with the KS statistics. The $p$ value was calculated as the fraction of the surrogate distributions with $KS$ values smaller than the $KS$ value of the corresponding experimental avalanche distribution. We took the significance level to be 0.05, i.e., for $p < 0.05$ the power law hypothesis was rejected, whereas for $p \geq 0.05$ the power law hypothesis was not rejected. To test whether average avalanche size scaled with duration according to $< S > \sim D^{\beta}$, we estimated the fitted $\beta$ from the experimental data using linear regression. We then compared the fitted $\beta$ to the predicted $\beta = (\alpha - 1)/(\tau - 1)$. We defined the absolute difference between fitted $\beta$ and predicted $\beta$ as the "Deviation from Criticality Coefficient" (DCC) and adopted this number as a useful measure of the deviation of the network state from criticality.

**Model Investigation**

We simulated a model network consisting of 4750 excitatory and 250 inhibitory binary probabilistic model neurons with sparse connectivity and external inputs. Connectivity between excitatory neurons and from excitatory to inhibitory neurons was 3%. Connectivity from inhibitory to excitatory neurons was varied between 1 and 60%. The strength of the connection from neuron $j$ to neuron $i$ is quantified in terms of the transition probability $P_{ij}$, which is the probability that a spike in neuron $j$ causes a spike in neuron $i$ in the next simulation time step[8]. In order to allow for inhibitory connections, $P_{ij}$ was allowed to be negative for these connections. The binary state $X_i(t)$ of neuron $i$ denotes whether the model neuron spikes ($X_i(t) = 1$) or does not spike ($X_i(t) = 0$) at time $t$. At each time step, the state of all neurons was updated synchronously according to the following update rule:

$$X_i(t+1) = \Theta\left[(1-\eta_i(t))\sum_j P_{ij}X_j(t) + \eta_i(t) - \xi_i(t)\right]$$

where $\xi_i(t)$ is a random number in [0 1] drawn from a uniform distribution, and $\Theta$ is the step function. The external input $\eta_i(t)$ quantifies the probability of that neuron spiking due to the external input alone. External input was added to 10 percent of the excitatory neurons and a variable percentage (20, 40, and 80%) of the inhibitory neurons. The external input was chosen to be smaller than the transition probability $P_{ij}$, which itself was small for large networks, $P_{ij} \sim 1/N$, where *N* is the total number of neurons in the network. Because of the weak external inputs, we can employ the approximation $1 - \eta_i \approx 1$ in the above update rule. The external input $\eta_i(t)$ was modeled as a binary Poisson process followed by smoothing with a Gaussian filter with a width of 20 time steps. The maximum eigenvalue $\lambda$ of the transition probability matrix $P_{ij}$ describes the network state: $\lambda < 1$ denotes subcritical regime, $\lambda \approx 1$ denotes the near critical regime and $\lambda > 1$ denotes the supercritical regime[8]. Here, $\lambda = 1.02$ corresponded to networks operating at the critical regime. Because of the sparse connectivity, most transition probability matrix elements are zero. For the existing connections, the $P_{ij}$ values were drawn from a uniform distribution and then scaled by a constant factor to reach the desired maximum eigenvalue $\lambda$. By default, $P_{ij}$ was constant in the first part of the simulation.

After the first 20,000 time steps, two forms of synaptic dynamics were permitted. Spike-timing dependent plasticity (STDP) was added according to:

$$\delta P_{ij}^{EE}(t) = K_{STDP}\left[X_i^E(t)X_j^E(t-1) - X_j^E(t)X_i^E(t-1)\right]$$

$$\delta P_{ij}^{EI}(t) = -K_{inh}X_j^I(t-1)[1 - X_i^E(t)\left(1 + \frac{1}{\mu_I}\right)]$$

where $\delta P_{ij}^{EE}(t)$ is the change of connection strength from excitatory neuron to excitatory neuron while $\delta P_{ij}^{EI}$ is the change of connection strength from inhibitory neuron to excitatory neuron induced by STDP and $\mu_I$ is the mean firing rate of all I neurons during baseline[9]. The parameter $K_{STDP}$ and $K_{inh}$ is a constant gain value. Synaptic scaling (SS) was added to both E and I cells according to $\delta P_{ij}^{SS}(t) = K_{SS}[F_i^{ref} - F_i^{hist}(t)]$, where $\delta P_{ij}^{SS}(t)$ is the change in synaptic strength according to SS and the parameter $F_i^{ref}$ is the reference firing rate for each single neuron, which provides a set point of firing rate, and $F_i^{hist}(t)$ is the weighted firing rate across a 15,000 step time window. In other words, SS was a global, multiplicative change in all of the synaptic inputs onto a neuron that functioned to compensate for alterations in the neuron's output. SS in excitatory neurons affected both excitatory and inhibitory inputs, while SS in inhibitory neurons affected only inhibitory to inhibitory connections. We found that firing rate and network dynamics are relatively stable in the presence of STDP and SS. To mimic a homeostatic challenge in the model, we reduced the external input $\eta_i(t)$ in an exponential manner leveling out at half the starting value.

Models that satisfied the following constraints were considered successful: 1) the firing rates of both excitatory and inhibitory neurons dropped below a threshold of 75% of normalized baseline activity, 2) the absolute value of the difference of the maximum eigenvalue of the transition probability matrix and 1.02 was larger than 0.05 before the dropping of the excitatory neuron firing rate, i.e. the network became non-critical prior to the disruption of excitatory neuronal firing rate. This corresponds to a DCC of >0.2 (Fig 1C). At the end of the experiment, the sum of the excitatory and inhibitory neuron firing rate difference from baseline firing rate was less than 0.09. i.e. firing rates rebounded to baseline levels (**Fig S3**, epsilon). Likewise, at the end of the experiment, the maximum eigenvalue returned to a near-critical zone (i.e. within 0.05 of 1.02, defined *a priori* as criticality in this context). This corresponds to a DCC to a value of <0.2 (Fig 1C).
